# Met is required for oligodendrocyte progenitor cell migration in *Danio rerio*

**DOI:** 10.1101/2021.05.21.445204

**Authors:** Maria F. Ali, Andrew J. Latimer, Yinxue Wang, Leah Hogenmiller, Laura Fontenas, Adam J. Isabella, Cecilia B. Moens, Guoqiang Yu, Sarah Kucenas

## Abstract

During vertebrate central nervous system development, most oligodendrocyte progenitor cells (OPCs) are specified in the ventral spinal cord and must migrate throughout the neural tube until they become evenly distributed, occupying non-overlapping domains. While this process of developmental OPC migration is well characterized, the nature of the molecular mediators that govern it remain largely unknown. Here, using zebrafish as a model, we demonstrate that Met signaling is required for initial developmental migration of OPCs, and, using cell-specific knock-down of Met signaling, show that Met acts cell-autonomously in OPCs. Taken together, these findings demonstrate *in vivo,* the role of Met signaling in OPC migration and provide new insight into how OPC migration is regulated during development.

## Introduction

During vertebrate spinal cord development, OPCs are specified from ventral gliogenic precursors (Lu et al. 2000; Dimou et al. 2008; Ravanelli et al. 2018). Immediately following specification, these cells undergo a process termed tiling where they actively disperse throughout the spinal cord, ultimately forming non-overlapping domains with neighboring OPCs (Kirby et al. 2006; De Biase et al. 2017). Although tiling is a behavior that has been extensively studied in the context of neuronal development, very few studies have characterized these events in glia, although this behavior is commonly reported in the literature (Cameron and Rao 2010; Grueber and Sagasti 2010; Villar-Cerviño et al. 2013; López-Hidalgo et al. 2016; De Biase et al. 2017; Nichols et al. 2018). The process of OPC tiling is comprised of four main cellular behaviors: migration, proliferation, cell death, and contact-mediated repulsion (Kirby et al. 2006; Hughes et al. 2013; Huang et al. 2020). Though the phenomenon of OPC tiling is well-described, the molecular mediators of this process remain largely unknown. What has been identified are a number of chemoattractant or chemorepellent molecules that influence OPC migration (Spassky et al. 2002; Tsai et al. 2002; Jarjour et al. 2003; Chu et al. 2017). In this study, we sought to identify, in an unbiased manner, molecular mediators that govern the initial migration of OPCs during developmental tiling. Using an unbiased, small molecule screen, we identified several pathways, including Met signaling, as essential mediators of OPC migration.

Previously, *in vitro* studies revealed that hepatocyte growth factor (Hgf), the ligand for Met, acts as a mitogenic signal for OPCs (Lalive et al. 2005; Ohya et al. 2007) Met, also known as scatter factor receptor or Hgf receptor, is a widely studied receptor tyrosine kinase that is involved in a number of morphogenetic processes during embryogenesis, including regulating cellular migration and motility (Soriano et al. 1995; Birchmeier and Gherardi 1998; Prat et al. 1998; Viticchiè et al. 2015; Zhang and Babic 2015). In particular, OPCs in culture, upon application of Hgf, exhibit increased migration and proliferation (Lalive et al. 2005; Ohya et al. 2007). Additionally, it is well documented that OPCs express the c-Met receptor (Kilpatrick et al. 2000; Lalive et al. 2005; Ohya et al. 2007; Mela and Goldman 2013). Though these studies established foundational work supporting Met as a possible mediator of OPC migration, further investigation into the role of Met signaling in regulating developmental OPC migration *in vivo* were impeded because mouse *Met* mutants are embryonic lethal.

In order to study the role of Met signaling *in vivo,* we utilized zebrafish as a vertebrate model. Because zebrafish embryos receive maternal mRNAs, including *met* mRNA from their mother (Latimer and Jessen 2008), zebrafish *met* mutants successfully complete embryogenesis and can therefore be used to investigate later developmental processes, including OPC migration. In fact, many recent studies have used zebrafish embryos and larvae lacking Met function to study a number of development processes including motor axon targeting and migratory muscle precursor migration (Nord et al. 2019; Talbot et al. 2019; Isabella et al. 2020).

Here, we describe the identification of the Met receptor as an essential mediator of OPC migration. Using a combination of pharmacological and genetic manipulation with *in vivo,* time-lapse imaging and a new software to analyze migration dynamics, we demonstrate that Met signaling is required for the initial, dorsal migration of OPCs during development of the vertebrate spinal cord. Furthermore, by utilizing cell-specific dominant negative Met (DNmet) constructs, we show that Met acts cell-autonomously within OPCs. Together, our results demonstrate that Met signaling regulates initial OPC migration during developmental tiling.

## Materials and Methods

### Zebrafish Husbandry

All animal studies were approved by the University of Virginia Institutional Animal Care and Use Committee. Zebrafish strains used in this study were: AB*, *met^uva38^, met^fh534^, Tg(sox10(4.9):tagrfp)^uva5^,Tg(olig2:egfp)^vu12^* (Shin et al. 2003)*; Tg(sox10(7.2):mrfp)^vu234^* (Kucenas et al. 2008)*; met^egfp^*, *Tg(sox10(4.9):DNmet::IRES:egfp; cry:egfp)^uva40^; Tg(olig1(5.3):DNmet::IRES:egfp; cry:egfp)^uva39^*; and *hgfa^fh529^* (Isabella et al. 2020). Table 1 denotes abbreviations used for each mutant and transgenic line. Embryos were raised at 28.5°C in egg water and staged by hours or days post fertilization (hpf and dpf, respectively) (Kimmel et al. 1995). Embryos of either sex were used for all experiments. Phenyl-thiourea (PTU) (0.004%) (Sigma) in egg water was used to reduce pigmentation for imaging. Stable, germline transgenic lines were used in all experiments.

**Table 1.** Zebrafish lines used in this study and their genotypes.

| Full Name | Abbreviation | Reference |
| --- | --- | --- |
| AB* | wildtype |  |
| *Tg(olig2:egfp)*<sup>vu12</sup> | *olig2:egfp* | Shin et al. 2003 |
| *Tg(sox10(4.9):tagrfp)*<sup>uva5</sup> | *sox10:tagrfp* | Zhu et al. 2019 |
| *Met*<sup>fh555Tg</sup> | *met*<sup>egfp</sup> | This paper |
| *Tg(sox10(7.2):mrfp)*<sup>vu234</sup> | *sox10:mrfp* | Kucenas et al. 2008 |
| *met*<sup>uva38</sup> | *met*<sup>uva38/uva38</sup> | This paper |
| *Tg(sox10(4.9):DNmet::IRES:egfp; cry:egfp)*<sup>uva40</sup> | *sox10:DNmet* | This paper |
| *Tg(olig1(5.3):DNmet::IRES:EGFP; cry:EGFP)*<sup>uva39</sup> | *olig1:DNmet* | This paper |
| *hgfa*<sup>fh529</sup> | *hgfa*<sup>fh529/fh529</sup> | Isabella et al. 2020 |
| *met*<sup>fh534</sup> | *met*<sup>fh534/fh534</sup> | This paper |

### Generation of transgenic lines

All constructs were generated using the Tol2kit Gateway-based cloning system (Kwan et al. 2007). Vectors used for making the expression constructs were p5E-sox10(-4.9) (Carney et al. 2006), p5E-olig1(-5.3) (this paper), pME-DNmet (this paper), p3E-IRES-EGFPpA, and pDestTol2pA2 destination vector (Kwan et al. 2007).

To generate p5E-olig1, we amplified 5.3 kb of sequence immediately upstream of the olig1 gene (Ensembl: ENSDARG00000040948) from wildtype genomic DNA using the following primers: forward 5’-GTATGAAGCCTCTTGGCACAG-3’ and reverse 5’-CTGAAAAAAGATATTCAGAGAACATGG-3’, as previously described (Auer et al. 2018). The resulting PCR product was subcloned into pENTR™ 5’-TOPO (Invitrogen) to generate a p5E entry for Gateway cloning and was verified by sequencing.

To generate pME-DNmet, we used the Q5 Site-Directed Mutagenesis Kit (NEB) and generated site-specific mutations in *met* cDNA, which we generated used RT-PCR as described below, using the following primers: forward 5’-CAACATCGACAAAATGACACCCTTCCCCTCTCTCATATCATCTCAG-3’ and reverse, 5’-ACGAAGGTGGTGTTCAGGAGGATGAAGTGCTCTCCGCTGAAGC-3’. We confirmed the mutations using sequencing, then subcloned the cDNA containing the DNmet mutations into pME-MCS to generate a pME entry for Gateway cloning. p5E, pME, and p3E-IRES-EGFPpA vectors were ligated into destination vectors through LR reactions (Ashton et al. 2012). Final constructs were amplified, verified by restriction digest, and sequenced to confirm the insertions. To generate stable transgenic lines, plasmid DNAs were microinjected at a concentration of 25 ng/μL in combination with 10 ng/μL *Tol2* transposase mRNA at the one-cell stage and screened for founders (Kawakami 2004).

The *met^fh555Tg^* enhancer trap line was generated using the CRISPR/Cas9-mediated knock-in strategy described in (Kimura et al. 2014). 1 nL of a cocktail of the following components was injected into one-cell stage embryos: 66.6 ng/μL each of 3 CRISPR guide RNAs targeting within or just upstream of the *met* 5’ UTR (target sequences GGTCTCGGGATGGGATGCGA, GGTTCTCTCCGCAAACGCTG, and GGGTAAGCGGGTTCGCTGAT), 200 ng/μL Mbait gRNA, 20 ng/μL Mbait-hsp70-GFP plasmid, 800 ng/μL Cas9 protein (PNABio #CP02). Founders were screened for *GFP* expression replicating the known *met* expression pattern.

The *met^fh534^* allele was made using standard Crispr/Cas9 mutagenesis protocol (Talbot and Amacher 2014). One cell stage embryos were injected with 100 pg each of 2 CRISPR guide RNAs (target sequences GGTTCTGGCCATCTGGCTCG and GGCTTCGGCTGCGTGTTTCA) and 500 pg Cas9 protein, and F1 mutant animals were identified by sequencing. *met^fh534^* is a 25bp deletion starting at nucleotide 3275 of the coding sequence, resulting in a frameshift at amino acid 1093 and a premature stop at amino acid 1105.

### *In vivo* imaging

Embryos were anesthetized with 0.01% 3-aminobenzoic acid ester (Tricaine), immersed in 0.8% low-melting point agarose and mounted in glass-bottomed 35 mm Petri dishes (Electron Microscopy Sciences). After mounting, the Petri dish was filled with egg water containing PTU and Tricaine. A 40X water objective (NA = 1.1) mounted on a motorized Zeis AxioObserver Z1 microscope equipped with a Quorum WaveFX-X1 (Quorum Technologies) or Andor CSU-W1 (Oxford Instruments) spinning disc confocal system was used to capture all images. Images were imported into MetaMorph (Molecular Devices) and/or ImageJ. Time-lapse images were analyzed using cell tracking software as previously described in (Wang et al. 2018). All images were then imported in Adobe Photoshop and Illustrator. Adjustments were limited to levels, contrast, and cropping.

### Cell tracking software

For quantification and motility analysis of OPCs, we developed an automated software to detect and track motile cells in time-lapse imaging experiments of *olig2:egfp* embryos and larvae (Wang et al. 2018). To correct for photobleaching, we normalized fluorescence intensity to an identical mean and an identical variance at all time points and to account for long-term image shift due to larval growth, we used global image registration. For intra-frame detection and segmentation of all cells, we applied the cell detection algorithm SynQuant to map the second-order derivative transformed intensity (Wang et al. 2020). The under-segmentation and over-segmentation in intra-frame detection were then corrected by imposing temporal consistency of segmentation, which was modeled as an optimization problem. Rapid motion was detected by testing regional intensity change and to link the detections, we adapted our established algorithm muSSP to a mixed motion model form (Wang et al. 2019).

The motion patterns of all obtained traces were then analyzed to obtain a quantification of cell motility and to distinguish OPCs from other cells in the field of view. The distance traveled was calculated by adding up the magnitude of displacement between any two consecutive time points and the instantaneous velocity of any *olig2^+^* OPC was approximated by the average velocity between two consecutive time points, while the instantaneous speed was the magnitude of it. The overall average speed was the distance traveled divided by time period and we identified OPCs from neighboring cells by hypothesis testing on the instantaneous velocity. Assuming the majority of cells in the field of view were not moving, the null distribution was learned by fitting a multivariate Gaussian distribution to all instantaneous velocity values of the obtained traces. Any cell whose trace once had significantly high instantaneous velocity was identified as an OPC.

### Wholemount Immunohistochemistry

Dechorionated embryos and larvae were fixed with 4% PFA for 1 hour at room temperature (RT), washed in PBSTX (1% Triton X-100, 1x PBS) for 5 min, followed by a 5 minute wash with DWTx (1% Triton X-100 in DH_2_O), then permeabilized in acetone at RT for 5 min and at −20°C for 10 min, followed by a 5 min wash with PBSTx. Embryos were then blocked in 5% goat serum/PBSTx for 1 hr, incubated in primary antibody with 5% goat serum/PBSTx for 1 hr at RT and overnight at 4°C. Embryos were washed extensively with PBSTx at RT and incubated in secondary antibody overnight at 4°C. After antibody incubation, embryos were washed extensively with PBSTx and stored in PBS at 4°C until imaging. The following antibodies were used: rabbit anti-Sox10 (1:5000) (Binari et al. 2013), Alexa Fluor 647 goat anti-rabbit IgG(H+L) (1:1000; ThermoFisher). Embryos were mounted in glass-bottomed Petri dishes for imaging as described above.

### Immunohistochemistry on sections

Embryos and larvae were fixed in 4% PFA for 2 hr at RT. After fixation, the embryos were sectioned by embedding them in 1.5% agarose/30% sucrose and frozen in 2-methylbutane chilled by immersion in liquid nitrogen. We collected 20 μm transverse sections on microscope slides using a cryostat microtome. Sections were rehydrated in PBS for 1 hr and blocked with 5% goat serum/PBS for 1 hr at RT. Primary antibody incubation was done overnight at 4°C. Secondary antibody incubation was done for 2 hr at RT. Antibodies used were: rabbit anti-Sox10 (1:5000) (Binari et al. 2013), mouse 3D4 anti-met (1:100; ThermoFisher), Alexa Fluor 647 goat anti-rabbit IgG(H+L)(1:1000), and Alexa Fluor 568 goat anti-mouse IgG(H+L) (1:1000). Sections were covered with Aqua-Poly/Mount (Polysciences). A 63X oil objective (NA = 1.4) mounted on a motorized Zeiss AxioObserver Z1 microscope equipped with a Quorum WaveFX-X1(Quorum Technologies) or Andor CSU-W1 (Andor Oxford Instruments) spinning disc confocal system was used to capture all images. Images were imported into Image J and Adobe Photoshop and Illustrator. Adjustments were limited to levels, contrast, and cropping.

### Chemical treatments

In our initial small molecule screen, dechorionated *olig2:egfp* larvae were treated with 10 μm kinase inhibitor in 1% DMSO in PTU egg water from 24 to 76 hours post fertilization (hpf). Kinase inhibitors used were 1 of 430 kinase inhibitors from the L1200 Kinase Inhibitor Library (Selleck Chem), MK-2461 (Selleck Chem), or Trichostatin-A (TSA) (Selleck Chem). Control siblings were treated with 1% DMSO in PTU egg water. The small molecule screen was conducted in triplicate.

For the EdU incorporation assay, larvae were treated with 0.4 mM EdU in 4% DMSO from 70 to 74 hpf in PTU egg water at 28.5°C then fixed for 1 hr in 4% PFA at RT. Larvae were washed for 5 min with 1X PBSTX, 5 min in DWTX, then permeabilized with cold acetone for 10 min at −20 °C and stained for EdU using the Click-it EdU Cell Proliferation kit for Imaging with Alexa Fluor 647 dye (ThermoFisher), as detailed in the kit protocol. Click-it reaction was performed for 1 hr at RT and thoroughly washed overnight with PBSTX prior to imaging.

### Genotyping

Genomic DNA was extracted using HotSHOT (hot sodium hydroxide and tris) and PCR was performed using GoTaq green master mix (Promega) (Truett et al. 2000). The primers used for genotyping *met^uva38^* are as follows: forward 5’-ATCGTACGCATGTGTTCTTCAG-3’ and reverse 5’-TGATGTCCGTGATGGAGATAAG-3’. The primers used for genotyping *met^fh534^* are as follows: forward 5’-AATCTCTGCCATGTTTTCCTGT-3’ and reverse 5’-AGTCCAAAACTATCCCAAGCAA-3’.

### RT-PCR

mRNA was extracted and cDNA synthesized as described previously (Peterson and Freeman 2009) with the use of a RNA easy Mini kit (Qiagen) and High-capacity cDNA Reverse Transcription kit (Applied Biosystems) according to manufacturer instructions. Equal amounts of mRNA were used for cDNA synthesis, and PCR was performed using GoTaq green master mix (Promega). Primers used were: *ef1α* forward: 5’-GAGACTGGTGTCCTCAAGCC-3’ and reverse: 5’-CCAACGTTGTCACCAGGAGT-3’, and *met^uva38^* forward: 5’-ATCGTACGCATGTGTTCTTCAG-3’ and reverse: 5’-TGATGTCCGTGATGGAGATAAG-3’.

### Statistical Analysis

GraphPad PRISM 9 software was used to plot data and perform statistical analyses. Pairwise comparison *P-*values involving only 2 groups were calculated using a Student’s two-tailed *t*-test. Pairwise comparison *P-*values involving more than 2 groups were calculated using a one-way ANOVAs followed by Dunnett’s Multiple Comparison tests or Tukey’s Multiple Comparison test. The data in plots and the text are presented as means ± SEM.

## Results

### Met inhibition impairs developmental OPC migration

While the phenomena of OPC migration and tiling are well known, very few mediators of this process have been identified. Therefore, to identify molecular mediators of OPC tiling, we conducted an unbiased kinase inhibitor screen in zebrafish embryos and larvae. To do this, we treated *olig2:egfp* zebrafish embryos, where *olig2* regulatory sequences drive expression of GFP in motor neurons and oligodendrocyte lineage cells (OLCs), with 1% DMSO as a control, 0.2 μM Trichostatin A (TSA) in 1% DMSO as a control, or 10 μM of 1 of 430 kinase inhibitors from the Selleck Kinase Inhibitor Library in 1% DMSO. We used TSA as a positive control because it is a histone deacetylase (HDAC) inhibitor that blocks OPC specification and therefore, embryos treated with this compound would show significantly reduced OPC migration into the dorsal spinal cord (Cunliffe and Casaccia-Bonnefil 2006) (Figure 1A and 1B). We treated the embryos from 24 hours post fertilization (hpf), which is prior to OPC specification, until 76 hpf, which is during the middle-to-late migratory phase of these cells. By treating prior to OPC specification, we sought to identify molecular mediators that affect OPC migration, but do not block OPC specification. If we observed a complete absence of OPC migration, similar to that which occurs in the presence of TSA, then it is possible that the small molecule affected either OPC specification or migration. However, if we saw defects in migration, but there were still OPCs present, then we knew the kinase inhibitor likely did not affect specification. At 76 hpf, we first screened drug-treated larvae to confirm that overall larval morphology was indistinguishable compared to DMSO-treated larvae. We then individually screened drug-treated larvae for changes in OPC migration by looking for either an increase or decrease in the number of *olig2^+^* cells in the dorsal spinal cord compared to DMSO-treated controls (Figure 1B). From our screen of 430 kinase inhibitors, we identified 35 compounds that resulted in decreased numbers of OPCs in the dorsal spinal cord and 19 compounds that resulted in increased numbers of OPCs in the dorsal spinal cord. One exciting “hit” was in larvae treated with MK2461, a c-Met inhibitor, in which we observed a decrease in the number of OPCs in the dorsal spinal cord.

**Figure 1.**
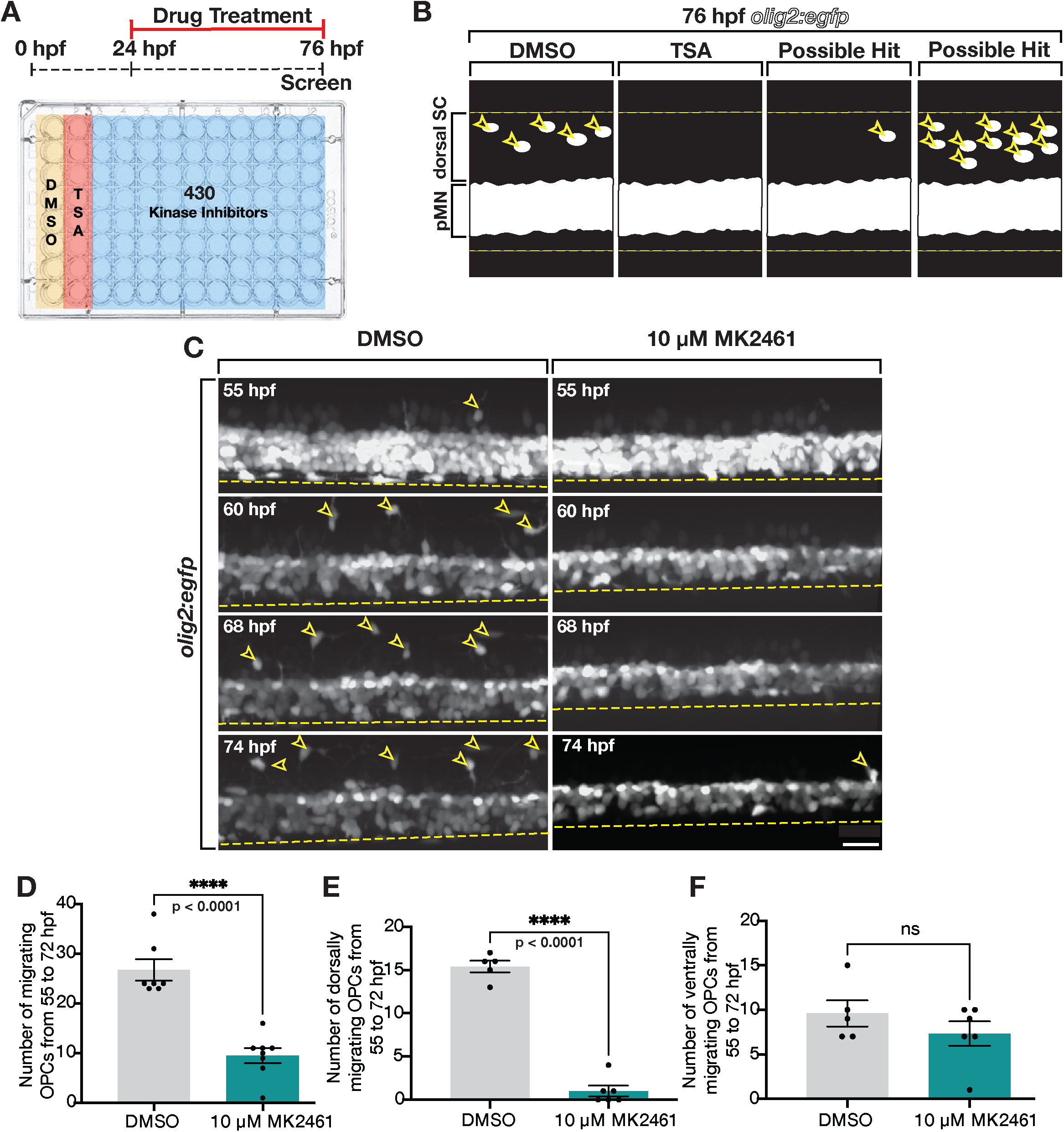
Kinase inhibitor screen identifies Met as mediator of dorsal OPC migration. (A) Schematic of kinase inhibitor screen and treatment paradigm that tested 430 kinase inhibitors for developmental OPC migration defects. Trichostatin A (TSA), which inhibits OPC specification, was used as a positive control. (B) Cartoon of a lateral view of 76 hpf *olig2:egfp* larvae spinal cord showing DMSO (negative control), TSA (positive control), and examples of possible hits: reduced OLCs in the dorsal spinal cord (SC) and increased OLCs in the dorsal SC. pMN denotes pMN domain. Yellow dashed lines mark the boundaries of the spinal cord and open yellow arrowheads mark dorsal OLCs. (C) Images taken from 18 hour time-lapse movies of DMSO and MK2461-treated 55 hpf *olig2:egfp* zebrafish larvae. Yellow open arrowheads denote dorsally migrating OPCs. Yellow dashed line denotes ventral edge of the spinal cord. (D-F) Quantifications taken from time-lapse movies of DMSO (n = 7) and MK2461-treated (n = 7) larvae in (C). Mean with SEM. Student’s t-test was used in D-F. Scale bar, 20 μm.

To confirm the reduction in dorsal OPCs we observed during MK2461 treatment, we used *in vivo,* time-lapse imaging in 55 hpf *olig2:egfp* larvae treated with MK2461 from 24 hpf on (Figure 1C). In these movies, we see a significant reduction in the number of dorsally migrating OPCs (Figure 1C). However, we did observe active OPC migration in the ventral spinal cord, indicating that OPCs are specified but exhibit migration defects (Figure 1C). For quantification and motility analyses of OPCs, we developed an automated software to detect and track motile *olig2^+^* OPCs distinct from *olig2^+^* cells in pMN domain of *olig2:egfp* larvae, which contains a mixture of migratory OPCs and non-migratory motor neurons cell and precursors (Movie 1) (Wang et al. 2018). Using this cell tracking software, we assessed the behaviors of OPCs in time-lapse movies from 55 to 74 hpf. From these analyses, we found that the number of migratory OPCs was significantly reduced in MK2461-treated larvae compared to control larvae (Figure 1D, Movies 2 and 3). Specifically, we observed significantly reduced numbers of dorsally, but not ventrally, migrating OPCs in MK2461-treated larvae compared to DMSO-treated controls in a 3-somite window (Figure 1E and 1F).

We next wanted to determine if treatment with MK2461 altered other aspects of OPC migration, including velocity and distance traveled. To do this, we used our cell-tracking software to analyze our time-lapse movies, like those shown in Figure 1C. Overall, MK2461-treatment did not alter velocity or distance traveled when looking at all OPCs compared to control larvae (Figure S1A and B). However, the dorsally migrating, and not ventrally migrating, OPCs traveled a shorter distance in MK2461-treated larvae compared to DMSO-treated control larvae (Figure S1C and E). The average velocity of migration for dorsally and ventrally migrating OPCs was not affected by MK2461 treatment (Figure S1D and F). Taken together, we observed that treatment with MK2461 resulted in a reduction in the number of OPCs that migrate dorsally during development and therefore, the Met receptor, is a likely mediator of OPC dynamics during development.

### Zebrafish OPCs express the Met receptor

Previous *in vitro* studies of mouse and rat OPCs used antibody labeling to show that OPCs express the Met receptor (Kilpatrick et al. 2000; Lalive et al. 2005; Ohya et al. 2007; Mela and Goldman 2013). Therefore, we wanted to determine if zebrafish OPCs also expressed Met. To investigate Met expression in OLCs, we used the c-Met 3D4 antibody to label met^+^ cells in conjunction with an antibody specific to zebrafish sox10 to label 3 days post fertilization (3 dpf) *olig2:egfp* zebrafish larvae. We then imaged transverse sections through the spinal cord of antibody-labeled larvae and observed met^+^/sox10^+^/*olig2^+^* cells in the spinal cord (Figure 2A). Interestingly, not all sox10^+^/*olig2^+^* cells were met^+^, indicating that there are populations of both met^+^ and met^-^OLCs (Figure 2A).

**Figure 2.**
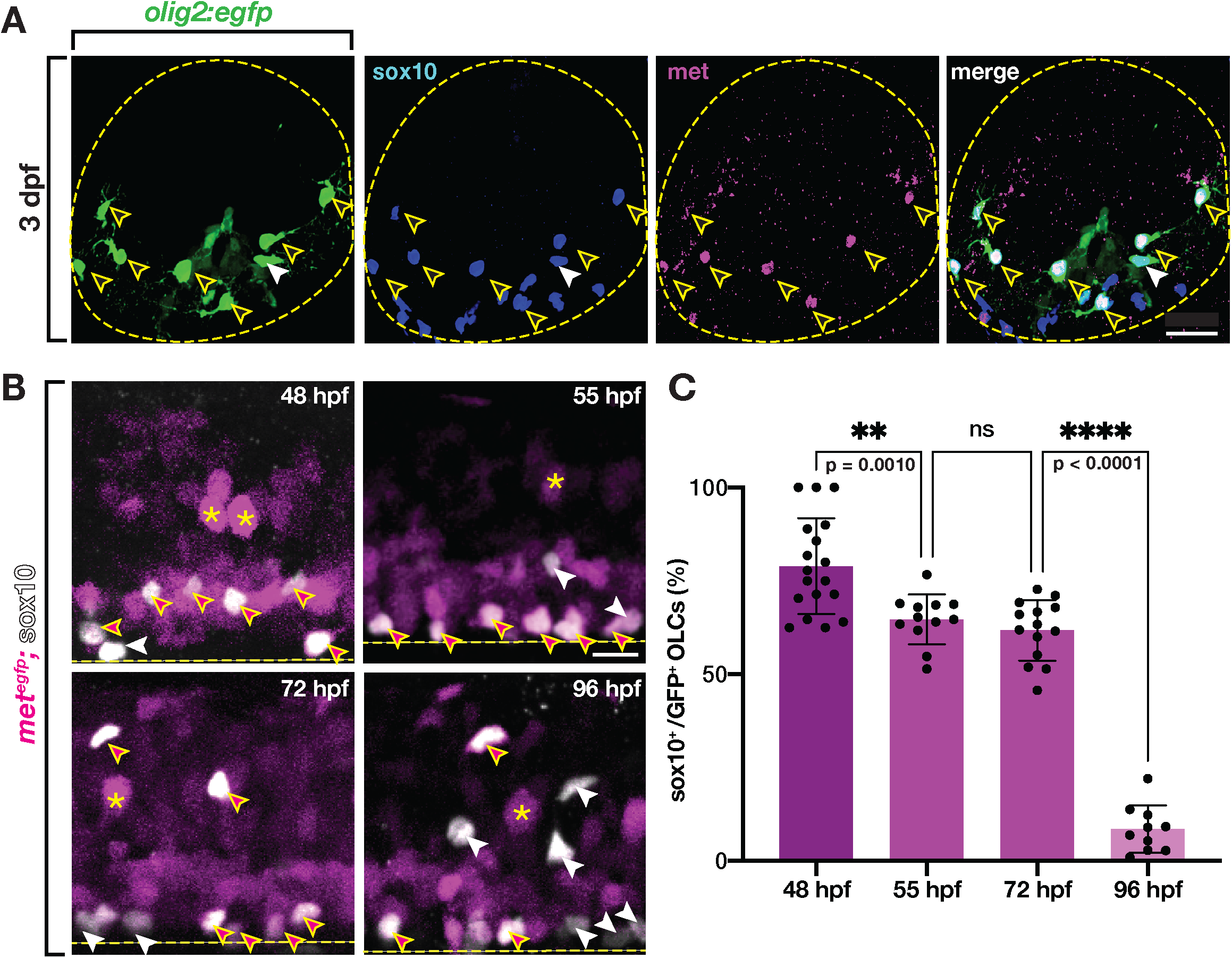
Zebrafish OLCs express Met. (A) Transverse section of a 3 dpf *olig2:egfp* zebrafish spinal cord labeled with antibodies to sox10 (blue) and met (magenta). Yellow open arrowheads denote sox10^+^/met^+^ OLCs. White arrowheads denote sox10^+^/met^-^ OLCs. Yellow dashed circle denotes boundary of the spinal cord. (B) Lateral view of *met^egfp^* zebrafish spinal cords at 48, 55, 72, and 96 hpf labeled with a sox10 antibody. Asterisks denote examples of *met^+^* motor neurons. Magenta-filled yellow arrowheads denote sox10^+^/*met^+^* OPCs, white arrowheads denote sox10^+^/*met^-^* OLCs. Yellow dashed line denotes ventral edge of the spinal cord. (C) Percentage of OLCs that are sox10^+^ and *met:gfp^+^* in 322 μm of spinal cord at 48 hpf (n = 18), 55 hpf (n = 12), 72 hpf (n = 14), and 96 hpf (n = 10). Mean with SEM. Statistical test: 1-way ANOVA with Tukey’s Multiple Comparison Test. Scale bars, 10 μm (A), 20 μm (B).

To confirm these findings, we used a new enhancer trap transgenic line, *met^egfp^,* where eGFP is under *met* regulation due to CRISPR-targeted insertion immediately upstream of the endogenous *met* gene, to assess *met* expression in OLCs from 48 to 96 hpf (Kimura et al. 2014). To label OLCs, we labeled *met^egfp^* embryos and larvae with our sox10 antibody (Figure 2B). We then imaged the spinal cord and quantified the number of *met^+^/* sox10^+^ cells. We found that at 48 hpf, just prior to OPC migration, a large percentage of sox10^+^ OPCs were *met^+^* (78.95%) (Figure 2C). By 55 hpf, only 64.68% of sox10^+^ OLCs were *met^+^* and this level of *met^+^* OLCs stayed roughly constant through 72 hpf, where 61.76% were *met^+^* (Figure 2C). The consistent expression of *met* in the majority of OLCs from 55 to 72 hpf is consistent with the major migratory period of OPCs, which occurs during this same time window. By 96 hpf, when most OLCs have completed their migration and many are initiating myelination of spinal cord axons, the population of OLCs that were *met^+^* was decreased significantly to 8.56% (Figure 2C). These findings are consistent with previous investigations demonstrating that OPCs down-regulate expression of Met in order to differentiate into oligodendrocytes (Ohya et al. 2007). From these expression studies, we conclude that zebrafish OLCs express Met.

Because we observed Met expression in migratory OPCs, we next sought to more closely investigate the effect of MK2461 on their migration. First, we conducted a dose response curve using the same treatment paradigm as the drug screen described above by treating 24 hpf *olig2:egfp* zebrafish larvae with a 1% DMSO control or increasing doses of MK2461 in 1% DMSO and quantified the number of *olig2^+^* OLCs in the dorsal spinal cord at 76 hpf. We found that increasing doses of MK2461 resulted in decreasing numbers of OLCs in the dorsal spinal cord compared to DMSO-treated larvae (p < 0.0001) (Figure 3A). To more directly assay the positioning and number of OLCs in control and drug-treated larvae, we used serial sectioning in 76 hpf *olig2:egfp;sox10:mrfp* zebrafish larvae treated with 1% DMSO or 10 μM MK2461 in 1% DMSO and quantified the number and location of *olig2^+^/sox10^+^* cells (Figure 3B). These studies revealed that the overall number of OLCs was reduced in the spinal cord of MK2461-treated larvae when compared to controls (Figure 3C), and the decrease affected both the number of dorsal and ventral OLCs (p < 0.0001) (Figure 3D and 3E). Interestingly, in these studies, we saw an increase in the number of OLCs in the pMN domain in larvae treated with MK2461, which indicates that OLCs were specified in MK2461-treated larvae and that the migration defect we observed was not simply due to perturbed specification (p = 0.0349) (Figure 3F). Additionally, in contrast to our earlier studies in whole larvae where we observed that ventral OLCs were unaffected by MK2461 treatment, we saw a significant reduction in this population when we assessed their location in serial sections (p < 0.0001) (Figure 3E). We believe this occurred because it is difficult to observe individual ventral OLCs in whole-mount *olig2*:*egfp* larvae. Finally, the overall of reduction of OLCs in MK2461-treated larvae could be the result of reduced proliferation in OPCs, as Met is also implicated in regulating OPC proliferation in *in vitro* studies using OPC cell culture (Ohya et al. 2007), and we investigate this possibility later in this manuscript. Taken together, we conclude that Met mediates OPC migration and potentially OPC proliferation during development.

**Figure 3.**
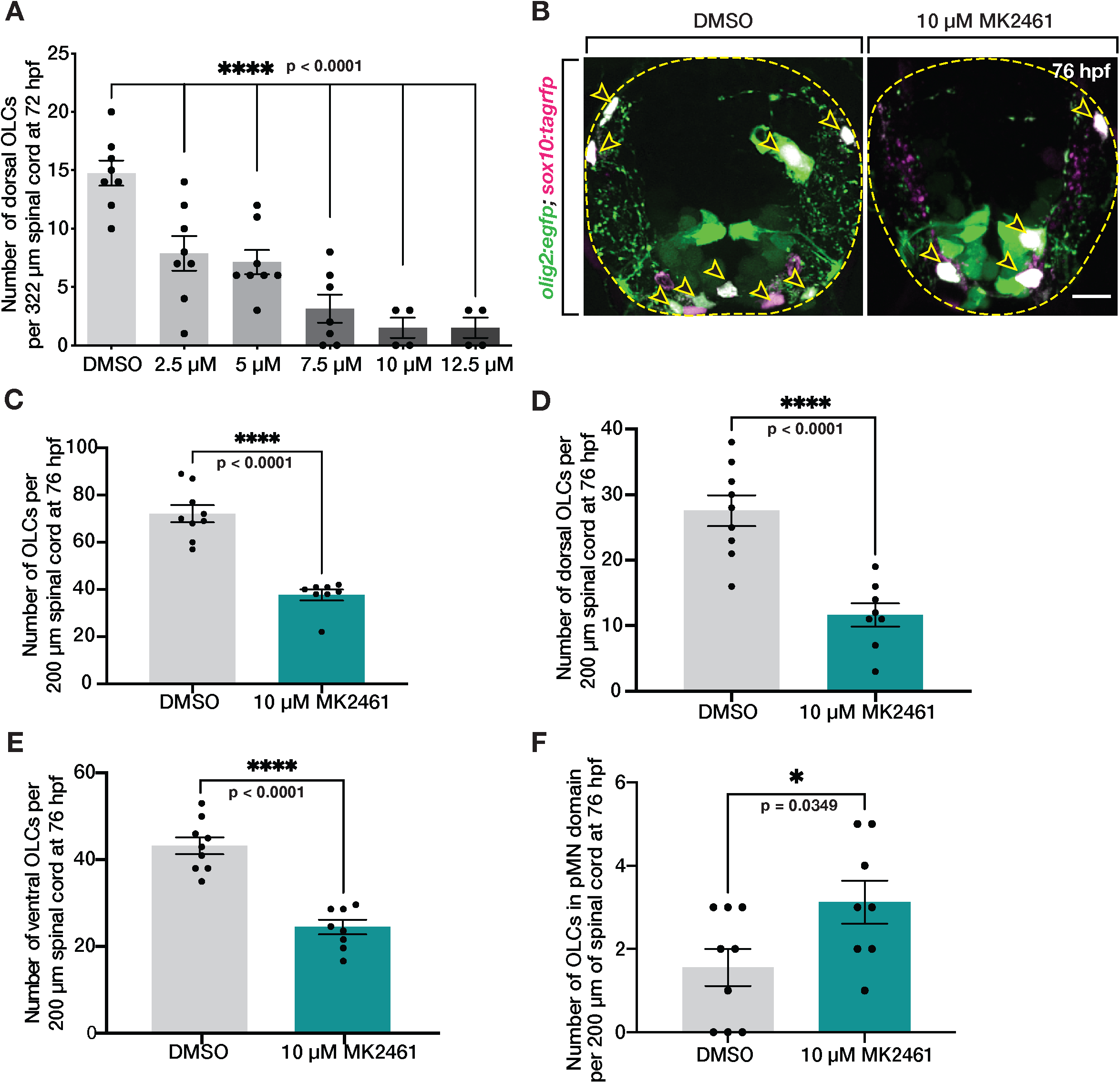
Met receptor inhibition decreases the number of OLCs in spinal cord. (A) Dose-response curve of the number of OLCs in the dorsal spinal cord of fish treated from 24 hpf to 3 dpf with 1% DMSO (n = 8) and 10 μM MK2461 in 1% DMSO in the following doses: 2.5 μM (n = 8), 5 μM (n = 8), 7.5 μM (n = 7), 10 μM (n = 4), and 12.5 μM (n = 8). Statistical test: 1-way ANOVA with Dunnett’s Multiple Comparison Test. (B) Transverse sections of 76 hpf *olig2:egfp;sox10:tagrfp* larvae treated with DMSO or 10 μM MK2461 from 24 hpf to 3 dpf. Yellow open arrowheads denote *sox10^+^/olig2^+^* OLCs. (C-F) Quantifications of *olig2^+^/sox10^+^* OLCs from serial sections of *olig2:egfp;sox10:tagrfp* larvae treated with DMSO (n = 9) or MK2461 (n = 8) from 24 hpf to 3 dpf. Mean with SEM. Student’s T-test was used in C-F. Scale bar, 10 μm.

### Met is required for initial developmental OPC migration

Given the migration phenotype we observed with the c-Met inhibitor, MK2461, and the expression pattern of Met in zebrafish OLCs, we sought to further investigate Met as a mediator of OPC migration during development using CRISPR/Cas9 mutagenesis (Hruscha et al. 2013; Hwang et al. 2013). Using CHOPCHOP, we generated a gRNA specific to exon 2 of the zebrafish *met* coding sequence (Labun et al. 2019). Using this synthesized gRNA, we injected one-cell embryos with the gRNA for *met* and Cas9 protein and grew up potential founders to adulthood. We then outcrossed putative founders and screened for frameshift mutations in their offspring and identified a founder with a mutation that resulted in a 16 base pair insertion in the second exon of *met* that results in an early stop codon (Figure 4A). This mutation causes the premature termination in the beta chain of the sema domain resulting in a truncated polypeptide that would be functionally unable to homodimerize upon Hgf binding and therefore, would be unable to initiate downstream signaling (Figure 4B). Importantly, these mutant larvae had an overall normal morphology and body length when compared to wildtype siblings (Figure S2A). To confirm the incorporation of a 16 base pair insertion, we isolated RNA from *met^+/+^, met^+/uva38^,* and *met^uva38/uva38^* larvae and performed RT-PCR. As expected, *met^uva38^* heterozygous and homozygous larvae had a larger band compared to wildtype that was consistent with a 16 bp insertion (Figure S2B). Finally, to confirm our *met* allele contained a deleterious mutation in *met,* we crossed a heterozygous *met^uva38^* adult with a *met^fh534^* heterozygous adult and imaged the spinal cord of 72 hpf *olig2:egfp* mutant larvae (Figure S2C). In these trans-heterozygous *met^uva38/fh534^* larvae, we observed a significant decrease in the number of dorsal OLCs when compared to wildtype siblings (Figure S2D).

**Figure 4.**
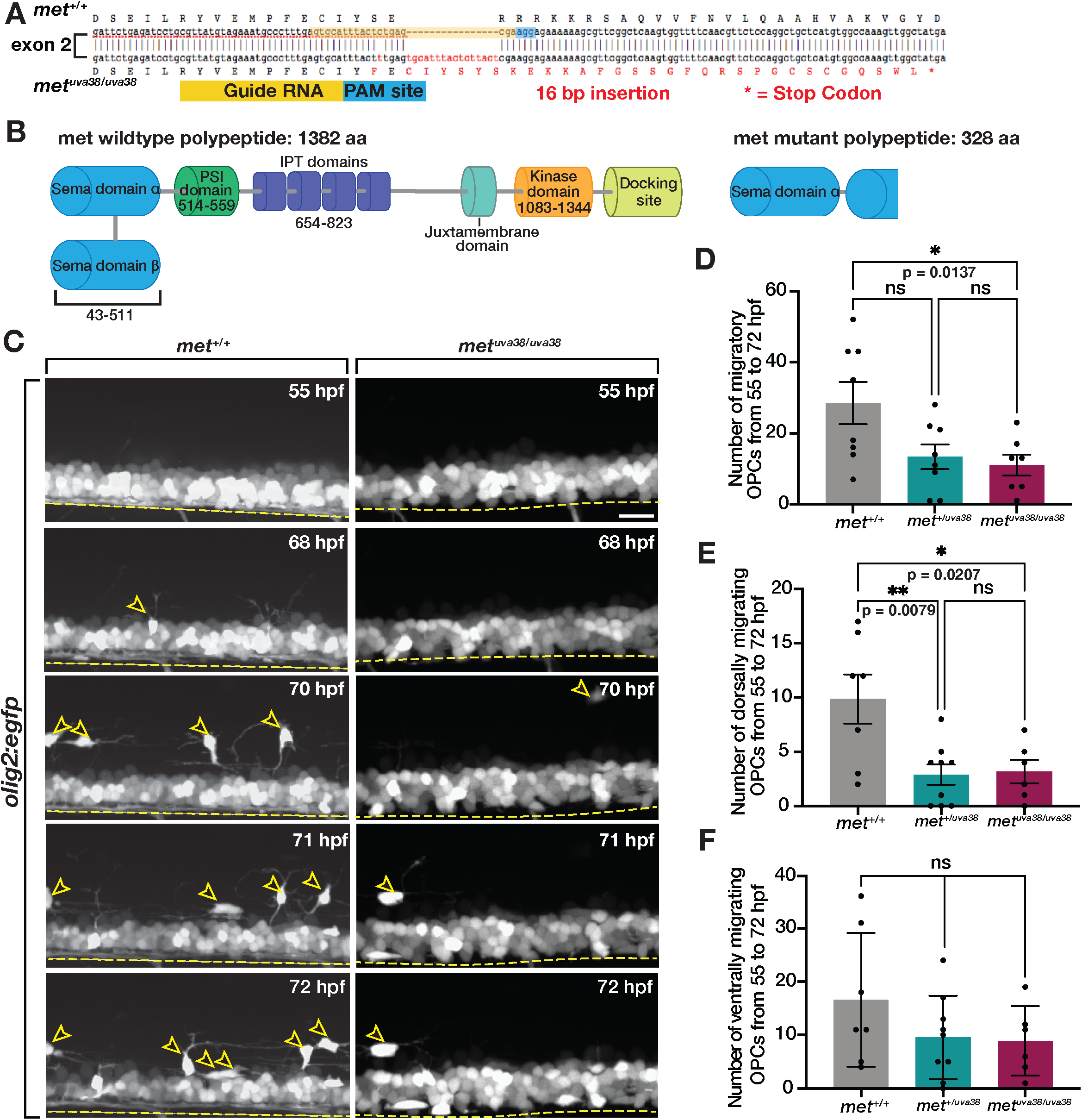
Met is required for initiation of dorsal OPC migration. (A) Diagram of *met* mutant created using CRISPR/Cas9 mutagenesis with gRNA target sequence (yellow) and PAM site (blue) resulting in a 16 base pair causing a frameshift mutation and early stop codon (asterisk). (B) Diagram of wildtype met protein and met mutant polypeptide sequences. (C) Images taken from 18 hour time-lapse imaging of 55 hpf *olig2:egfp met^+/+^,* and *met^uva38/uva38^* zebrafish larvae. Yellow open arrowheads denote dorsally migrating OPCs. Yellow dashed line denotes ventral edge of spinal cord. (D-F) Quantifications taken from time-lapse movies of 55 hpf *olig2:egfp met^+/+^* (n = 8), *met^+/uva38^* (n = 8), and *met^uva38/uva38^* (n = 6) zebrafish larvae. Mean with SEM. Statistical test: 1-way ANOVA with Tukey’s Multiple Comparison Test was used for D-F. Scale bar, 20 μm.

We next used this new *met^uva38^* allele to assess the role of *met* in regulating developmental OPC migration. Using *in vivo,* time-lapse imaging in combination with our cell-tracking software in 55 hpf *olig2:egfp;met^+/+^, met^+/uva38^,* and *met^uva38/uva38^* larvae, we analyzed OPC migration (Wang et al. 2018). This imaging of *olig2:egfp;met^uva38/uva38^* larvae revealed a significant decrease in the number of migrating OPCs compared to *met^+/+^* larvae (p = 0.0307) (Figure 4C and 4D). Interestingly, the decrease in OPC migration in the *met^uva38/uva38^* larvae primarily affected dorsally-migrating OLCs (p = 0.0207) (Figure 4E). There was also a decrease in dorsally migrating OPCs in *met^+/uva38^* larvae, which is likely due to the tightly regulated nature of Met signaling (p = 0.0079) (Figure 4E) (Zhang and Babic 2015). In contrast, ventrally migrating OPCs in *met^uva38/uva38^* larvae were similar to *met^+/+^* and *met^+/uva38^* larvae, although there was a trend of reduced numbers of ventral migratory OPCs (Figure 4F). Finally, consistent with our Met inhibitor studies, OPCs showed no difference in average velocity of migration or distance traveled when *met* was perturbed (Figure S3). Taken together, our results demonstrate that *met* is required for dorsal OPC migration during developmental OPC tiling.

To more closely examine the effect of a *met* mutation on developmental OPC migration, we performed serial sectioning on 76 hpf *olig2:egfp;sox10:mrfp;met^+/+^, met^+/uva38^,* and *met^uva38/uva38^* larvae. We then imaged transverse sections of the spinal cord and assessed the location and number of OLCs (Figure 5A). Similar to what we observed with the c-Met inhibitor, we observed a decrease in the overall number of OLCs in *met^uva38/uva38^* larvae compared to wildtype siblings (p = 0.0137) (Figure 5B). Additionally, we observed a reduction in the number of OLCs in both the dorsal and ventral portions of the spinal cord in *met^uva38/uva38^* larvae compared to their wildtype siblings (dorsal, p = 0.0335; ventral, p = 0.0014) (Figure 5C, 5D). This finding is slightly different than what we observed when looking at whole-mount, lateral views of the spinal cord in our *met* mutant larvae and larvae treated with MK2461. However, this difference is likely due to the fact that in lateral views, it is very difficult to visualize all ventral OLCs because of the expression of *olig2:egfp* in ventral spinal cord precursors and motor neurons. Therefore, our findings here with a careful analysis of OLC location in transverse sections is more accurate. Additionally, we observed an increase in the number of OLCs in the pMN domain in *met^uva38/uva38^* larvae compared to wildtype siblings, which is consistent with what we observed in larvae treated with MK2461 (p = 0.0013) (Figure 5E). This data also fits with what we observed in time-lapse movies of *met* mutant larvae, where we observed OPCs in the pMN domain of the spinal cord extend cellular processes into the dorsal spinal cord but then fail to migrate dorsally (Movies 4 and 5). The increased number of OPCs in the pMN domain in *met^uva38/uva38^* larvae compared to wildtype siblings, and the OPC process extension behavior in our time-lapse imaging indicate that there is a reduction in the number of OPCs that are migrating out of the pMN domain. This data supports our hypothesis that OPCs require *met* for initial migration out of the pMN domain during development, while sparing OPC specification.

**Figure 5.**
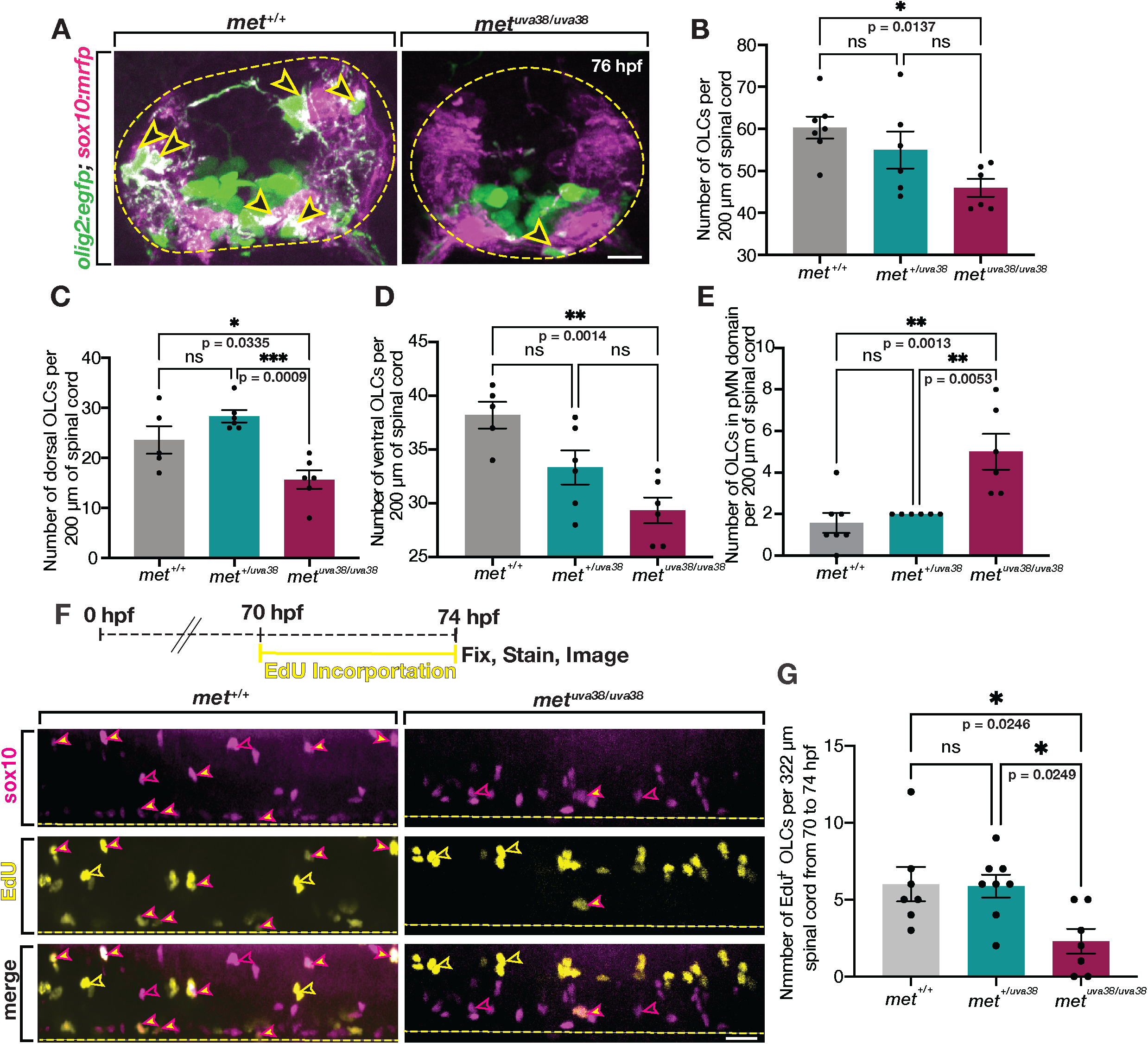
Met mutants exhibit reduced OPC proliferation. (A) Transverse sections of 76 hpf *olig2:egfp;sox10:mrfp met^+/+^* and *met^uva38/uva38^* larvae. Open yellow arrowheads denote *sox10^+^/olig2^+^* OPCs. Dashed yellow circle denotes boundary of the spinal cord. (B-E) Quantifications of *olig2^+^/sox10^+^* OLCs from serial sections of 76 hpf *olig2:egfp;sox10:mrfp met^+/+^* (n = 7), *met^+/uva38^* (n = 6), and *met^uva38/uva38^* (n = 6) larvae. Mean with SEM. Statistical test: 1-way ANOVA with Tukey’s Multiple Comparison Test. (F) EdU treatment paradigm and lateral view spinal cord images of 74 hpf *olig2:egfp;sox10:mrfp met^+/+^* and *met^uva38/uva38^* zebrafish larvae. Magenta-outlined yellow arrowheads denote *sox10^+^/olig2^+^/*EdU^+^ OPCs. Magenta open arrowheads denote *sox10^+^/olig2^+^/*EdU^-^ OLCs. Yellow open arrowheads denote *sox10^-^/olig2^-^*/EdU^+^ cells. Yellow dashed line denotes ventral edge of the spinal cord. (G) Quantifications of *sox10^+^/olig2^+^/*EdU^+^ OLCs from spinal cord images of 76 hpf *olig2:egfp;sox10:mrfp* EdU labeled *met^+/+^* (n = 7), *met^+/uva38^* (n = 8), and *met^uva38/uva38^* (n = 7) larvae. Mean with SEM. Statistical test: 1-way ANOVA with Tukey’s Multiple Comparison Test. Scale bars, (A) 10 μm, (F) 20 μm.

Because Hgf is the ligand for the Met receptor, we wanted to investigate whether there would also be a reduction in dorsal OLC numbers in *hgfa* mutants as well (Isabella et al. 2020). To do this, we used sox10 antibody labeling on 72 hpf *hgfa* mutants and imaged transverse sections of the spinal cord to quantify the number of sox10^+^ OLCs (Figure S4A). We hypothesized that we would see a similar change in OPC migration to what we observed in *met* mutant larvae. Excitingly, we observed a decrease in the overall number of OLCs in *hgfa^fh529/fh529^* larvae compared to wildtype siblings (p = 0.0008) (Figure S4B). Furthermore, we found that there was a significant decrease in the number of dorsal OPCs in *hgfa^fh529/fh529^* larvae compared to wildtype siblings (p < 0.0001), though the number of ventral OLCs was not significantly different compared to wildtype siblings (Figure S4C and D). Finally, we saw an increase in the number of OLCs in the pMN domain in *hgfa^fh529/fh529^* larvae compared to wildtype siblings (p = 0.0004) (Figure S4E). These data are all consistent with what we observed in our *met* mutant larvae and further support that Met signaling is required for developmental OPC migration.

From these data, we hypothesize that the reduction in the number of OLCs in *met^uva38/uva38^* larvae could be caused by either a reduction in OPC specification, which would lead to fewer OLCs in the spinal cord, or a reduction in OPC proliferation, which would result in wildtype numbers of OPCs during specification, but fewer OPCs during the migratory period. Met has been implicated in regulating cellular proliferation in a number of different cell types including hepatocytes, melanocytes, and other epithelial cell types (Tamagnone and Comoglio 1997; Prat et al. 1998; Viticchiè et al. 2015). To investigate if the overall decrease of OPCs in *met^uva38/uva38^* larvae is a consequence of reduced OPC specification, we did serial sectioning of 48 hpf *olig2:egfp;sox10:mrfp;met^+/+^, met^+/uva38^,* and *met^uva38/uva38^* embryos (Figure S5). We chose 48 hpf because it is sufficiently after the window of OPC specification which begins at 36 hpf, but prior to the main migratory period of OPCs which begins at approximately 55 hpf (Kirby et al. 2006; Appel et al. 2018). In these studies, we found that there was no difference in the number of OPCs in *met^+/+^, met^+/uva38^,* and *met^uva38/uva38^* larvae at 48 hpf, which demonstrates that OPC specification is not affected by *met* mutation (Figure S5).

With OPC specification being unaffected in *met^uva38/uva38^* larvae, we next hypothesized that the reduction in the number of OLCs in *met^uva38/uva38^* larvae was caused by a decrease in OPC proliferation, which has previously been demonstrated (Birchmeier and Gherardi 1998; Ohya et al. 2007; Viticchiè et al. 2015). To investigate OPC proliferation, we treated *olig2:egfp;sox10:mrfp;met^+/+^, met^+/uva38^,* and *met^uva38/uva38^* larvae with EdU (5-ethynyl-2’-deoxyuridine) from 70 to 74 hpf in order to detect DNA synthesis in proliferative cells (Figure 5F). We then imaged spinal cords of EdU-treated larvae and quantified *olig2^+^/sox10^+^/*EdU^+^ OLCs (Figure 5F). We observed that *met^uva38/uva38^* larvae had fewer EdU^+^ OLCs compared to wildtype and heterozygous siblings at 74 hpf (wt, p = 0.0246; heterozygous, p = 0.0249) (Figure 5G). These results demonstrate that the reduction in OLCs in *met^uva38/uva38^* larvae is due to a decrease in proliferation of OLCs and that *met* plays a role in regulating OLC proliferation and migration during development.

### Met inhibition in pre-migratory OPCs reduces dorsal migration out of the ventral spinal cord

During development, Met is expressed in a number of developing CNS neural populations, including motor neurons (Latimer and Jessen 2008). Because normal neuronal development influences OPC development (Appel et al. 2018), we wanted to ensure that the phenotypes we observed in our inhibitor-treated and whole animal *met* mutant studies were not due to non-cell autonomous effects on OPCs. Therefore, we created cell lineage specific, Met dominant negative constructs to specifically perturb met signaling in OLCs.

To do this, we used site-directed mutagenesis to selectively mutate three critical tyrosines in the docking domain of the c-Met receptor (Figure 6A), which, following Met dimerization, are trans-phosphorylated allowing for adaptor protein binding and downstream signaling (Figure 6B) (Soriano et al. 1995; Birchmeier and Gherardi 1998; Viticchiè et al. 2015). Previously published dominant negative Met (DNmet) constructs containing phenylalanine substitutions in the three docking site tyrosines of c-Met were successfully used in *in vitro* cell culture and zebrafish larvae for cell-specific reduction of Met signaling (Bardelli et al. 1999; Firon et al. 2000; Giordano et al. 2002; Latimer and Jessen 2008). The mutation of these tyrosines into phenylalanines physically prevents autophosphorylation of the docking site, thus preventing adaptor protein binding and downstream signaling (Ponzetto et al. 1996; Bardelli et al. 1999; Firon et al. 2000). We employed the same approach of creating point mutations that result in amino acid substitution of tyrosine to phenylalanine for the same critical tyrosines in the docking domain of the c-Met receptor as these previously published DNmet constructs (Firon et al. 2000; Giordano et al. 2002; Latimer and Jessen 2008). We confirmed successful mutation using Sanger sequencing, then drove our DNmet construct using either a *sox10* or *olig1* promoter (Figure 6D). The *sox10:DNmet* construct reduces Met signaling in glial cells and OPCs upon their specification beginning at approxiamtely 36 hpf (Figure 6C). The *olig1:DNmet* construct reduces Met signaling in OLCs at approximately 60 hpf, during their migratory phase (Auer et al. 2018) (Figure 6C). We additionally included an *IRES:GFP* reporter to more easily genotype and identify animals that contain the dominant negative constructs (Figure 6D). Taken together, we can use these constructs to temporally control Met signaling in OLCs by modulating it in either pre-migratory OPCs using *sox10:DNmet,* or migratory OLCs using *olig1:DNmet*.

**Figure 6.**
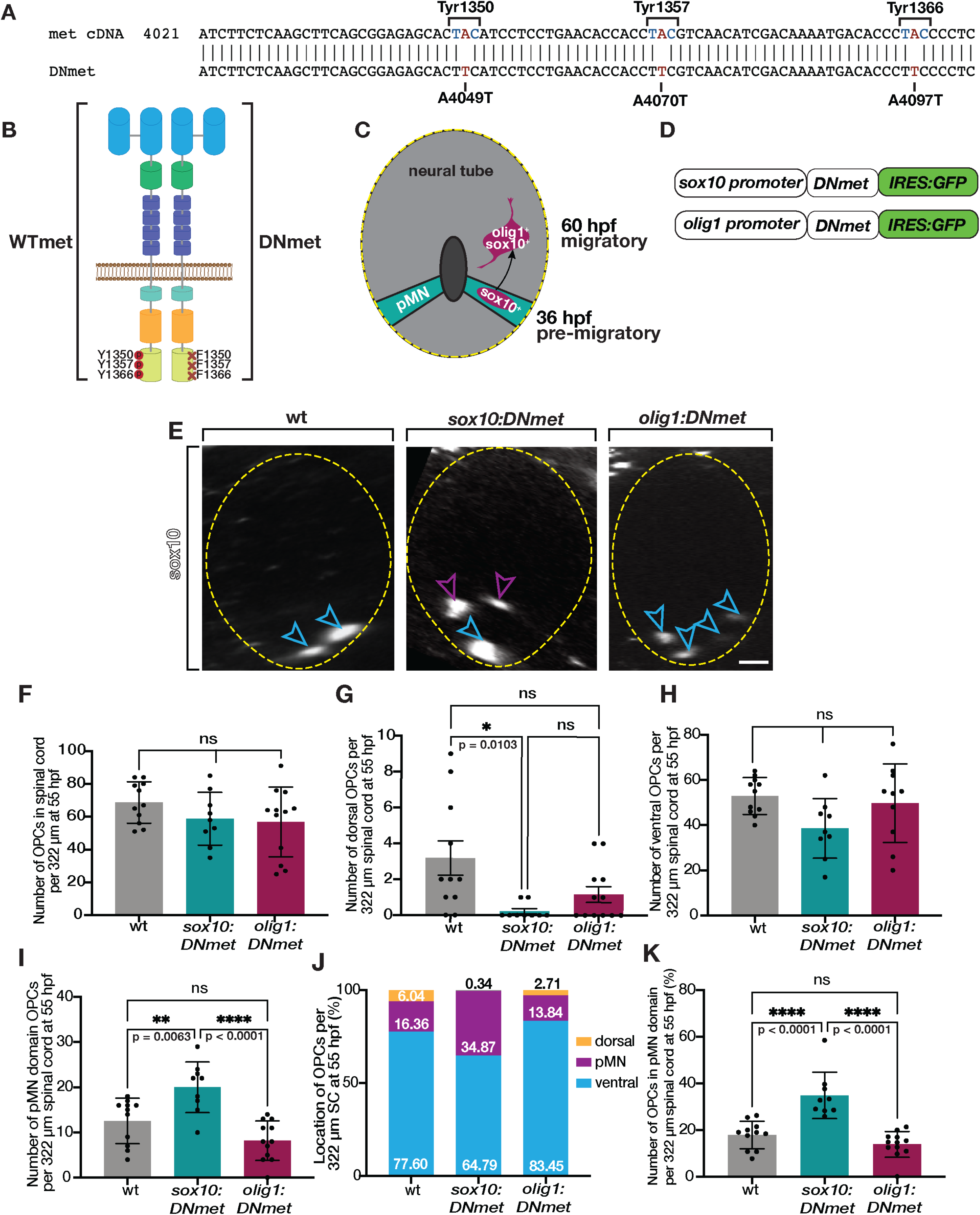
*sox10:DNmet* reduces OPC migration at 55 hpf. (A) Schematic of met receptor showing site-specific mutations converting adenines (A) to thymines (T) resulting in amino acid changes of tyrosines (Y) to phenylalanines (F). (B) Diagram of amino acid substitutions in docking site tyrosines created using site-directed mutagenesis. (C) Diagram of developing neural tube showing *sox10* turns on in pre-migratory OPCs around 36 hpf and *olig1* turns on in migratory OPCs around 60 hpf. (D) Schematic of DNmet constructs showing *DNmet* is driven by either a *sox10* or *olig1* promoter and includes *IRES:GFP* coding sequence. (E) Transverse sections of sox10 antibody labeled 55 hpf wildtype, *sox10:DNmet*, and *olig1:DNmet* larvae. Blue open arrowheads denote ventral OPCs. Purple open arrowheads denote pMN domain OPCs. Yellow dashed circle denotes spinal cord boundary. (F-K) Quantifications taken from images of 55 hpf sox10 antibody labeled wildtype (n = 10), *sox10DN:met* (n = 8), and *olig1:DNmet* (n = 12) zebrafish spinal cords. Mean with SEM. Statistical test: 1-way ANOVA with Tukeys’s Multiple Comparison Test. Scale bar, 20 μm.

To investigate the effect of OPC-specific reduction of Met signaling, we used sox10 antibody labeling to assess the position of OLCs in the spinal cord in *sox10:DNmet* and *olig1:DNmet* embryos and larvae (Figure 6E). At 55 hpf, we observed that the number of OPCs in both *sox10:DNmet* and *olig1:DNmet* larvae were unchanged compared to wildtype siblings (Figure 6F), supporting our conclusion that Met is not required for OPC specification. However, we observed decreased numbers of dorsal OPCs in *sox10:DNmet* larvae when compared to wildtype siblings at this stage (p = 0.0103) (Figure 6G). In contrast, while the number of ventral OPCs was unchanged compared to wildtype and *olig1:DNmet* larvae (Figure 6H), the *sox10:DNmet* larvae exhibited a significant increase in the number of OPCs in the pMN domain compared to both wildtype (p = 0.0063) and *olig1:DNmet* larvae at 55 hpf (p < 0.0001) (Figure 6I). These results demonstrate a significant reduction in dorsal migration of OPCs out of the pMN domain when Met signaling is reduced in pre-migratory OPCs using the *sox10:DNmet* construct. Additionally, while the overall positioning of the OPCs in the spinal cord among the three groups at 55 hpf demonstrated a large population of ventral OPCs, *sox10:DNmet* larvae exhibited an expanded population of pMN domain OPCs that was significantly different from both wildtype (p < 0.0001) and *olig1:DNmet* larvae (p < 0.0001) (Figure 6J and 6K). These results demonstrate that inhibition of Met signaling in pre-migratory OPCs causes a significant shift in the distribution of the position of OPCs toward the pMN domain at 55 hpf, indicating that OPCs require Met signaling to migrate dorsally during development.

We wanted to further examine the effect of reduced Met signaling in OLCs by looking at OLC positioning in the spinal cord toward the end of the migratory period at 72 hpf. We used sox10 antibody labeling to identify OLCs in the spinal cord and found that, at 72 hpf, the number of OLCs remained unchanged compared to wildtype in both the *sox10:DNmet* and *olig1:DNmet* zebrafish larvae (Figure 7A and 7B). Interestingly, at 72 hpf, the number of dorsal OLCs was reduced in *sox10:DNmet* larvae compared to both wildtype (p = 0.0042) and *olig1:DNmet* (p = 0.0461) larvae (Figure 7C). Additionally, ventral OLCs were reduced in *sox10:DNmet* larvae compared to wildtype (p = 0.0286) (Figure 7D). Concordantly, the number of pMN domain OLCs in *sox10:DNmet* was significantly increased compared to both wildtype (p < 0.0001) and *olig1:DNmet* (p < 0.0001) larvae (Figure 7E). Furthermore, the significantly increased percentage of OLCs in the pMN domain in *sox10:DNmet* larvae resulted in a reduced percentage of both dorsal and ventral OLCs in *sox10:DNmet* larvae compared to both wildtype and *olig1:DNmet* larvae at this stage (Figure 7F and 7G). Additionally, *olig1:DNmet* larvae had a slightly increased percentage of pMN domain OLCs when compared to wildtype, though significantly less than *sox10:DNmet* (p < 0.0001), indicating that reducing Met signaling later in developmental can also affect OLC positioning (p = 0.0147) (Figure 7G). Taken together, these data demonstrate that reducing Met signaling specifically in pre-migratory OPCs causes a reduction in dorsal OPC migration following specification. Additionally, this reduction in migration is not observed when Met signaling is knocked down in OLCs that are already in the migratory window. Therefore, Met acts cell-autonomously to induce OPC migration, following specification, during developmental OPC tiling.

**Figure 7.**
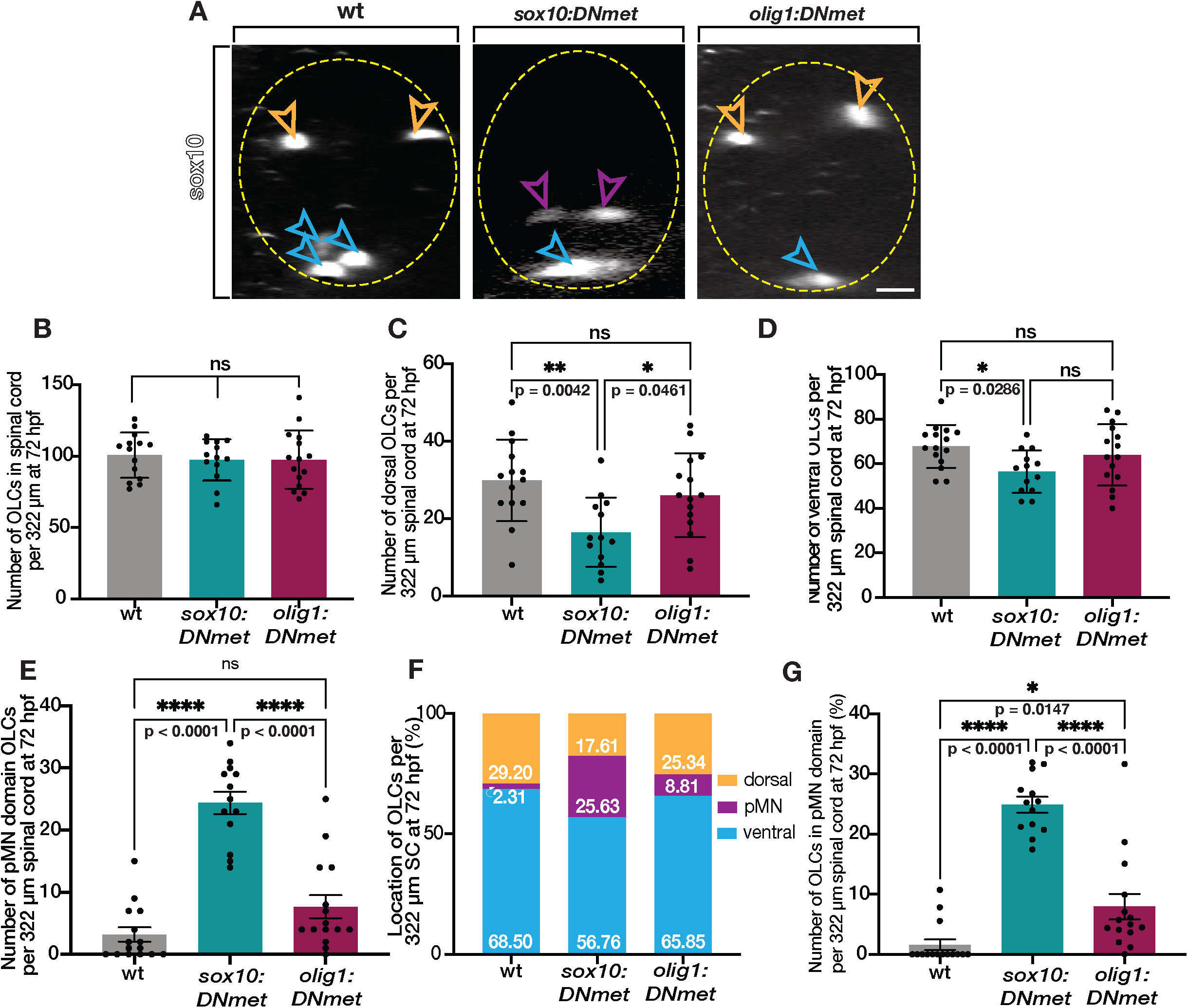
Met knock-down migration defects in pre-migratory OPCs persist to 72 hpf. (A) Transverse sections of sox10 antibody labeled 72 hpf wildtype, *sox10:DNmet*, and *olig1:DNmet* zebrafish larvae. Orange open arrowheads denote dorsal OLCs. Purple open arrowheads denote pMN domain OLCs. Blue open arrowheads denote ventral OLCs. Yellow dashed circle denotes spinal cord boundary. (B-G) Quantifications taken from images of 72 hpf sox10 antibody labeled wildtype (n = 14), *sox10DN:met* (n = 13), and *olig1:DNmet* (n = 15) zebrafish spinal cords. Mean with SEM. Statistical test: 1-way ANOVA with Tukey’s Multiple Comparison Test. Scale bar, 20 μm.

## Discussion

OPCs are a group of highly motile and dynamic cells that are specified during early development. Following specification, they migrate rapidly to become evenly distributed throughout the CNS in a process termed tiling. While this process is described phenomenologically in the literature, there is still much left to be discovered about how tiling is molecularly regulated (Kirby et al. 2006; Hughes et al. 2013). Using an unbiased small molecule screen and various genetic approaches, we sought to identify molecular mediators of the initial phase of OPC tiling, dorsal migration. In our studies, we showed that Met, the receptor for Hgf, is required for dorsal OPC migration and OLC proliferation. Additionally, using cell-specific reduction of Met signaling, we show that Met is required for initial migration in pre-migratory OPCs, specifically.

This work represents a significant advance in the understanding of Met signaling in OPC development because it is the first *in vivo* investigation demonstrating that loss of Met signaling reduces OPC migration. Previous *in vitro* studies demonstrated that applying Hgf to primary OPC cultures induced chemotaxis and proliferation (Kilpatrick et al. 2000; Lalive et al. 2005; Ohya et al. 2007; Mela and Goldman 2013). This *in vitro* data combined with our *in vivo* work demonstrates that Met expression on OPCs regulates their migration.

Interestingly, in our studies, we observed a trend in *met^+/uva38^* larvae where they occasionally exhibited OLC migration phenotypes similar to *met* mutants. A handful of studies have demonstrated that heterozygous *met* mutations can result in pathologically relevant outcomes (Ma et al. 2005; Matzke et al. 2007). More investigation into the regulation of Met signaling in OPCs and the normal physiological levels of active Met on OPCs would provide insight into the consequences of perturbing Met signaling in OPCs. Based on our observations here, it is likely that a certain threshold of Met signaling must be achieved for wildtype developmental processes to proceed and our heterozygous *met* mutants do not achieve this level of Met signaling resulting in the observed intermediate phenotypes.

What still remains unclear regarding the Met signaling pathway and OPC migration, is the source of Hgf. Some studies suggest that chemotropic factors released by astrocytes are influential in regulating OPC migration and positioning in the spinal cord (Tsai et al. 2002; Frost et al. 2009; Singh et al. 2019). Hgf/Met signaling could be functioning in a similar way. RNA-seq datasets show that astrocytes express high levels of Hgf during development (Zhang et al. 2014; Lake et al. 2016). Further investigations into Hgf expression and signaling in combination with newly published tools to investigate astrocytes in zebrafish could provide a wider lens into how Hgf/Met signaling regulates OPC migration (Chen et al. 2020).

Beyond developmental tiling, OPCs remain evenly tiled throughout life and utilize similar processes of migration, proliferation, contact-mediated repulsion, and apoptosis to maintain their spacing (Hughes et al. 2013; Tsata et al. 2019). While a number of chemotactic molecules have been investigated in developmental OPC tiling, adult tiling and how it is maintained remains largely unknown. Another benefit of using zebrafish is that homozygous *met* mutants can live into adulthood and therefore, can be utilized to study these tiling processes in mature animals (Nord et al. 2019). It is possible that, while adult OPCs remain in their distinct domain with little migration, Met could be utilized in response to injury. For example, following the loss of OLCs, surviving OPCs proliferate and migrate into OLC-depleted regions (Kirby et al. 2006; Kang et al. 2010; Hughes et al. 2013; Birey and Aguirre 2015). Additionally, previous studies using the disease model for multiple sclerosis in mice, experimental autoimmune encephalomyelitis (EAE), demonstrate that macrophages and microglia release Hgf and OPCs upregulate Met signaling in response to demyelinated legions (Lalive et al. 2005; Moransard et al. 2009). Beyond these studies, little is known about how OPCs respond to injury and initiate migration and it is possible that OPCs utilize the same migration mechanisms in both development and injury-response. With so much still unknown about how OPC behaviors are molecularly-mediated, this work demonstrates the critical role that Met plays in regulating initial OPC migration during development. Future work will need to be done to identify how Met signaling is regulated in OPCs and what role, if any, Met plays in other OPC process, such as adult tiling and response to injury.

## Data Availability

Fish lines and plasmids are available upon request. The authors affirm that all data necessary for confirming the conclusions of the article are present within the article, figures, and table.

## Acknowledgements

We thank Lori Tocke for zebrafish care and members of the Kucenas Lab for valuable discussions. This work was funded by NIH/National Institutes of Neurological Disorders and Stroke (NINDS): R01NS107525 (SK), F31NS108660 (MA), and R01NS109425 (CBM); by NIH/National Institute of General Medical Sciences: T32GM008715 (MA); by the NIH/Eunice Kennedy Shriver National Institute of Child Health and Human Development: F32HD096860 (AJI); and by NIH/National Institute of Mental Health: R01MH11504 (GY).

## Figure Legends

**Figure S1. Met inhibition alters the distance that dorsal OPCs migrate.** (A-F) Quantifications taken from time-lapse movies of DMSO (n = 7) and MK2461-treated (n = 7) larvae in (Figure 1C). Statistical test: Student’s t-test.

**Figure S2. Met mutants exhibit a 16 bp insertion and reduced dorsal OLCs.** (A) Bright-field images *of met^+/+^, met^+/uva38^*, and *met^uva38/uva38^* siblings at 3 dpf reveal no developmental delay in *met^uva38/uva38^* larvae. (B) RT-PCR of mRNA transcripts extracted from 48 hpf *met^+/+^, met^+/uva38^*, and *^metuva38/uva38^* embryos demonstrating 16 bp insertion in *met^+/uva38^* and *met^uva38/uva38^* RNA transcripts. (C) Images of 72 hpf *olig2:egfp met^+/+^* and *met^uva38/uva38^* larvae spinal cords. Yellow open arrowheads denote dorsal OLCs. Dashed yellow line denotes ventral edge of the spinal cord. (D) Quantifications taken from spinal cord images of *met^+/+^* (n = 3), *met^+/uva38^* (n = 6), *met^+/fh534^* (n= 7), and *met^uva38/fh534^* (n = 6) larvae. Mean with SEM. Statistical test: 1-way ANOVA with Tukey’s Multiple Comparison Test. Scale bars, 0.5 mm (A), 20 μm (C).

**Figure S3. Met mutants do not exhibit defects in OPC distance traveled or velocity.** (A-G) Quantifications taken from 18 hour time-lapse movies of 55 hpf *olig2:egfp met^+/+^* (n = 8), *met^+/uva38^* (n = 8), and *met^uva38/uva38^* (n = 6) larvae in (4C). Statistical test: 1-way ANOVA with Tukey’s Multiple Comparison Test.

**Figure S4. *hgfa* mutants exhibit reduced OPC numbers.** (A) Transverse sections of sox10 antibody labeled 72 hpf wildtype, *hgfa^+/+^,* and *hgfa^fh529/fh529^* larvae. Yellow open arrowheads denote sox10^+^ OLCs. Dashed yellow circle denotes boundary of spinal cord. (B-E) Quantifications taken from images of 72 hpf sox10 antibody labeled *hgfa^+/+^* (n = 14), *hgfa^+/fh529^* (n = 11), and *hgfa^fh529/fh529^* (n = 14) larval spinal cords. Mean with SEM. Statistical test: 1-way ANOVA with Tukey’s Multiple Comparison Test. Scale bar, 10 μm.

**Figure S5. Met mutants exhibit wildtype OPC specification.** (A) Transverse sections of 48 hpf *olig2:egfp;sox10:mrfp; met^+/+^* and *met^uva38/uva38^* embryos. Open yellow arrowheads denote *sox10^+^/olig2^+^* OPCs. Dashed yellow circle denotes boundary of spinal cord. (B-E) Quantifications of *olig2^+^/sox10^+^* OPCs from serial sections of 48 hpf *olig2:egfp;sox10:mrfp met^+/+^* (n = 6) and *met^+/uva38^* (n = 4), and *met^uva38/uva38^* (n = 4) embryos. Mean with SEM. Statistical test: 1-way ANOVA with Tukey’s Multiple Comparison Test. Scale bar, 10 μm.

**Movie 1. New cell tracking software labels migratory *olig2^+^* cells in *olig2:egfp* larvae from 55 to 76 hpf.** Images were taken every 10 minutes and the movie runs at 10 frames per second (fps).

**Movie 2. DMSO-treated *olig2:egfp* larvae exhibit wildtype OPC migration from 55 hpf to 76 hpf.** Images were taken every 10 minutes and the movie runs at 10 fps.

**Movie 3. MK2461-treated *olig2:egfp* larvae exhibit reduced dorsal OPC migration from 55 to 76 hpf.** Images were taken every 10 minutes and the movie runs at 10 fps.

**Movie 4. *olig2:egfp;met^+/+^* larvae exhibit wildtype OPC migration from 55 to 76 hpf.** Images were taken every 10 minutes and the movies runs at 10 fps.

**Movie 5. *olig2:egfp;met^uva38/uva38^* larvae exhibit reduced dorsal OPC migration from 55 to 76 hpf.** Images were taken every 10 minutes and the movie runs at 10 fps.

